# SPADE: Spatial Deconvolution for Domain Specific Cell-type Estimation

**DOI:** 10.1101/2023.04.14.536924

**Authors:** Yingying Lu, Qin Chen, Lingling An

## Abstract

The advent of spatial transcriptomics technology has allowed for the acquisition of gene expression profiles with multi-cellular resolution in a spatially resolved manner, presenting a new milestone in the field of genomics. However, the aggregate gene expression from heterogeneous cell types obtained by these technologies poses a significant challenge for a comprehensive delineation of cell type-specific spatial patterns. Here, we propose SPADE (SPAtial DEconvolution), an in-silico method designed to address this challenge by incorporating spatial patterns during cell type decomposition. SPADE utilizes a combination of single-cell RNA sequencing data, spatial location information, and histological information to computationally estimate the proportion of cell types present at each spatial location. In our study, we showcased the effectiveness of SPADE by conducting analyses on synthetic data. Our results indicated that SPADE was able to successfully identify cell type-specific spatial patterns that were not previously identified by existing deconvolution methods. Furthermore, we applied SPADE to a real-world dataset analyzing the developmental chicken heart, where we observed that SPADE was able to accurately capture the intricate processes of cellular differentiation and morphogenesis within the heart. Specifically, we were able to reliably estimate changes in cell type compositions over time, which is a critical aspect of understanding the underlying mechanisms of complex biological systems. These findings underscore the potential of SPADE as a valuable tool for analyzing complex biological systems and shedding light on their underlying mechanisms. Taken together, our results suggest that SPADE represents a significant advancement in the field of spatial transcriptomics, providing a powerful tool for characterizing complex spatial gene expression patterns in heterogeneous tissues.

## Introduction

Spatial transcriptomics is a cutting-edge technology [1] that has revolutionized the field of transcriptomics by allowing studies of gene expression with unprecedented detail. The ability to identify the specific location of gene expression within a tissue is a game-changer, as it opens up new avenues for understanding the complex interplay between gene expression and tissue architecture. By profiling the transcriptome at a high resolution in a spatial context, we can gain insights into the cellular heterogeneity that underlies normal tissue function or disease states [2]. This has enormous implications for addressing a wide range of biological questions, including the cellular basis for the brain function, and precision for treatment for heart disease or cancer. Spatial transcriptomics has applied for studying the immune system. Profiling the transcriptome of immune cells in different tissues has led to the gain of new insights into how the immune system responds to infection and disease. Such approach has the potential for the development of new immunotherapies against cancer and other diseases with spatial precision, essential for effective treatment with minimal non-specific side effect [3].

Current spatial transcriptomics technologies are limited in yielding cell type specific information within a tissue region, prohibiting the capture of complete gene expression patterns at single cell resolution in space [4]. For example, imaging-based ST methods are incapable of measuring a large number of genes, although these methods can provide detailed information at a single cell or subcellular level. Practically, this type of methods measures hundreds of genes based on the selection of probes from a gene list, making them less applicable for exploratory investigation at the level of transcriptome. In contrast, gene expression for each spatial location can be measured across the entire transcriptome using sequencing-based methods, at the sacrifice of single cell resolution [5]. Because of the differences in cell-type compositions between locations of tissues, the data from sequencing may be inconsistent from subsequent analyses. For example, when attempting to identify differentially expressed genes across multiple spatial locations, the observed gene expression variations may not be solely influenced by spatial location, but rather by differences in the proportions of cell types [6]. Hence, there is a growing need for methodologies that can accurately depict and describe the spatial patterns of gene expression variations, while also accounting for the specificity of individual cell types.

Single-cell RNA sequencing provides an insight into the functions and characteristics of individual cells [7]. This enables analysis of cellular heterogeneity, identification of uncommon cell types or subpopulations within a tissue and studying gene expression patterns for individual cells. However, single-cell RNA-Seq is unable to reveal how cells are physically arranged within a tissue, or how gene expression is influenced by cellular localization. Because the phenotype, metabolic state, and physiological function of a cell can be affected by its environment and cell-cell interaction, cell type deconvolution will provide an appealing tool for characterizing the intricate tissue architecture and scrutinizing the spatial localization of cell types.

In recent years, various algorithms have been developed to analyze cell type composition using gene expression and single-cell data. Methods such as CIBERSORTx [7], MuSiC [8], SCDC [9], and PREDE [10] have been meticulously tested for deconvoluting bulk RNA-Seq data. Although these methods can theoretically be applied to spatial transcriptomic data, they have several limitations that must be considered [5, 6]. First, unlike bulk RNA-Seq data, spatial genomics is composed of locations that may contain different portions of cell types, and the number of cells within a location can be small. Therefore, treating each location equally in bulk data analysis is not appropriate. Second, applying bulk RNA-Seq-related deconvolution techniques to spatial expression data can be time and memory-intensive due to the large number of spots in spatial data. Third, the unique property of spatial transcriptomic data compared to bulk RNA-Seq data is the relationship between locations and cell types. Locations that are closer together are more likely to have similar cell types [6]. Therefore, decomposing spatial transcriptomics at each spot requires a careful consideration of location similarities and spatial correlations. To address this need, several spatially resolved cell type deconvolution methods have been developed, including widely used SPOTlight [11], spatialDWLS [12], RCTD [13], and SpatialDecon [14]. These methods are specifically designed for spatial transcriptomics data and can accurately estimate cell type proportions at each location. Despite their effectiveness, these methods do not take into account the relationship between spatial information and histology. In other words, they do not consider the spatial dependency of gene expression and tissue structure. Because spatial structures can have a significant impact on biological functions, failing to account for them can be misleading.

To address this limitation, we introduce a reference based method, SPADE, for performing cell type deconvolution of spatially resolved transcriptomics data. The SPADE algorithm defines the cell type deconvolution task as a constrained nonlinear optimization problem that seeks to find the best estimated proportion to minimize the overall relative error between the true gene expression and estimated gene expression with restriction by non-negativity and sum-to-one constraints. Because spatial transcriptomic data exhibit distinctive features, such as a few cell types are associated with a specific locations, locations and cell types are related, neighboring locations show a similar pattern, we utilized a recently developed spatial domain detection algorithm [15] to account for the gene expression pattern, spatial location and histology information. To solve the unbalanced cell type between locations, we take advantage of lasso regression, which can perform automatic feature selection to decide the presence of particular cell types within each spot. Here we illustrate the benefit of SPADE through simulations by comparing to existing spatial deconvolution methods, and applications to public spatial transcriptomics studies in different area.

## Results

SPADE adopts a three-steps approach to estimate the cell type proportions within a spatial domain. Figure 1 shows a schematic overview of the SPADE methodology. The first step is to identify the spatial domains and domain-specific genes within a tissue. To accomplish this, SPADE uses spaGCN [15], a graph convolutional network designed specifically for spatial transcriptomics data. SpaGCN integrates gene expression, spatial location and histology data to identify the spatial domains and the genes that are specific to each domain. The second step in the SPADE approach is to determine the number of cell types present within each domain. To accomplish this, SPADE employs a Lasso regression algorithm. The Lasso regression algorithm uses the domain-specific gene expression data to identify the optimal number of cell types present within each domain. The identified number of cell types is used to perform deconvolution analysis in the next step. The final step of SPADE is to estimate cell type proportions for each spatial domain. To accomplish this, SPADE uses domain-specific and cell type-specific features. The domain-specific features consist of genes that are differentially expressed within each domain and are utilized to approximate the proportion of each cell type within the domain. The same principle applies to cell type-specific features, which are genes that are specific to each cell type and are employed to determine the contribution of each cell type to the gene expression profile within a given domain. The outcome of SPADE analysis is the calculated cell type proportions for every spatial location.

**Figure 1.**
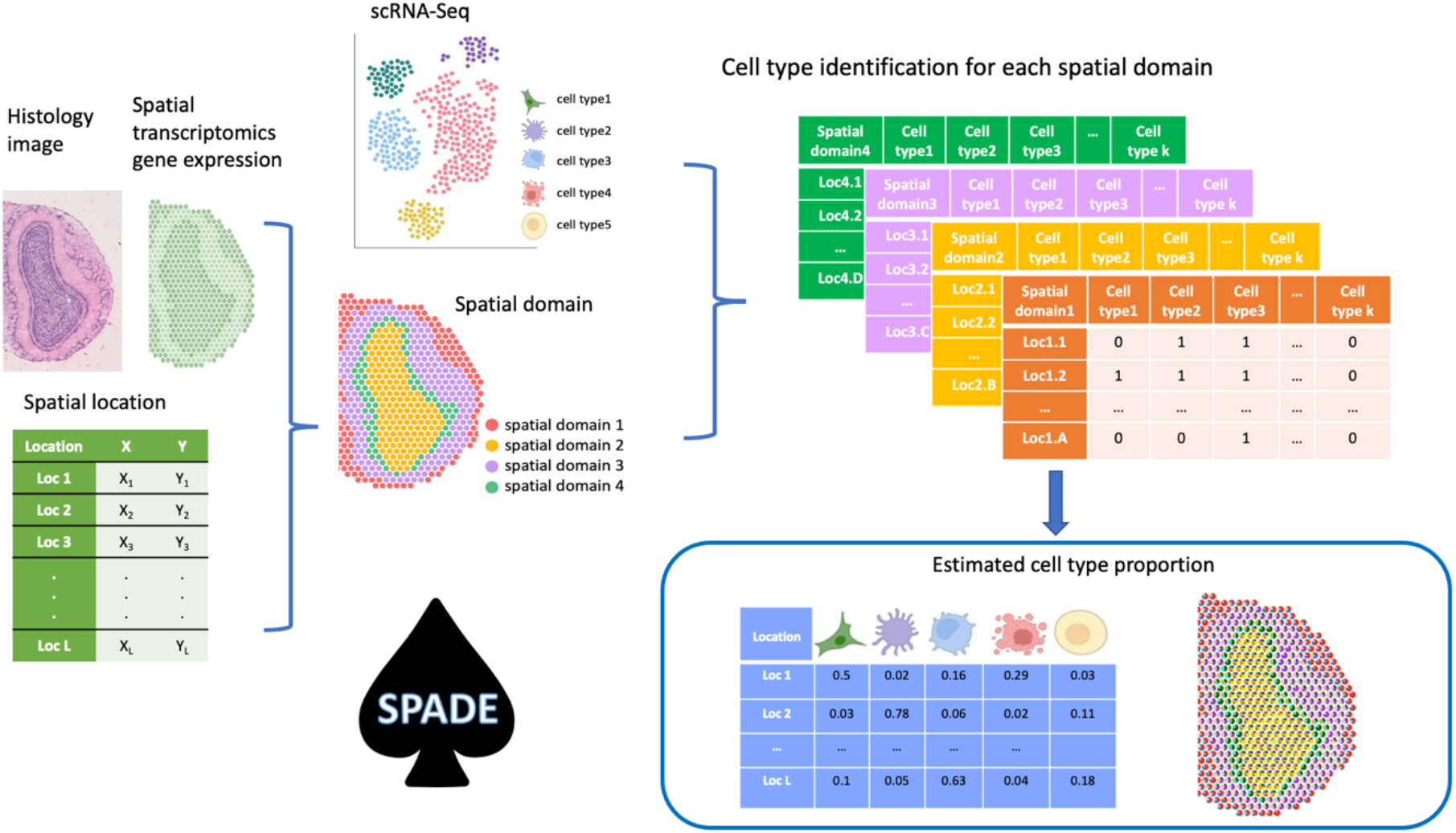
Schematic overview of SPADE. SPADE is a tool that designed for spatial transcriptomics data by uses reference single-cell RNA sequencing data to determine the cell type proportion at each location in the sample. It first uses histology, spatial location, and gene expression to identify spatial domains within a tissue. Subsequently, SPADE performs a comprehensive analysis for each domain by identifying the specific cell types present. Once the cell type information is obtained, SPADE utilizes scRNA-Seq data to perform deconvolution. The outcome of this process is the estimation of cell type proportions for every spatial location.

### Simulation setup

To generate synthetic spatial gene expression data, we followed the simulation idea of CARD [6] using the single-cell RNA-seq data. The generation of synthetic data was carried out in three stages. Firstly, we generated random proportions for each spatial location within each domain using a Dirichlet distribution. A dominant cell type was pre-defined, and the number of cell types per domain was limited to ensure cell type consistency across multiple replications. The remaining cell types for each domain were randomly selected based on the predetermined number. At the end of this stage, we obtained the true proportion and count matrix for cell types. Secondly, we selected cells from the single-cell RNA-seq data for each cell type based on the cell type count determined in stage one, and summed the counts to generate spatial transcriptomic data. Repeating this step\ yielded a gene-by-cell type matrix. Finally, we aggregated the cell type counts for each location for each gene to obtain the gene-by-location matrix, which serves as the pseudo spatial transcriptomic data.

We compared the method SPADE with existing spatial deconvolution methods, including CARD [6], SPOTlight [11], RCTD [13], spatialDWLS [12], and SpatialDecon [14]. CARD employs conditional autoregressive modeling that accounts for the spatial correlation structure across tissue locations. SPOTlight leverages both non-negative matrix factorization and non-negative least square to calculate cell type proportions in each spot. RCTD is based on cell type information generated from single-cell RNA-Seq data to deconvolute cell type compositions while counting for differences among sequencing technologies. SpatialDWLS extends DWLS and uses an enrichment test to determine cell types, then estimate cell type proportions by weighted least squares. SpatialDecon outperforms classic least-squares methods by log-normal regression and modeling backgrounds. We first run one simulation and compared the performance of the methods using three metrics: correlation, rooted mean square deviance, and mean absolute deviance. To test the robustness of SPADE, we then run 10 simulations and compared the total performance of all methods using the boxplot.

### Simulation using mouse olfactory bulb data

In the first study, we simulated spatial transcriptomic data using three publicly available datasets: a single-cell RNA-Seq dataset of the mouse olfactory bulb [16], a spatial gene expression dataset for the mouse olfactory bulb, and corresponding Hematoxylin and eosin stain (H&E) image data [17]. The integration of spatial gene expression dataset and image data enabled the extraction of domain information and spatial location. The results were illustrated in Figure 2a, which depicts a boxplot of all 10 replications, revealing that SPADE consistently outperformed other methods with the lowest mean absolute deviation (mAD), root mean squared error (RMSE), and the highest correlation across all simulations. To evaluate the ability of SPADE to detect the correct cell types in each spatial domain, in comparison with all other methods, we calculated true positive and false positive rates for each domain. The effectiveness of cell type selection in our lasso model was verified by the bar plot in Figure 2b, which indicates that SPADE correctly detected cell types for all domains.

**Figure 2:**
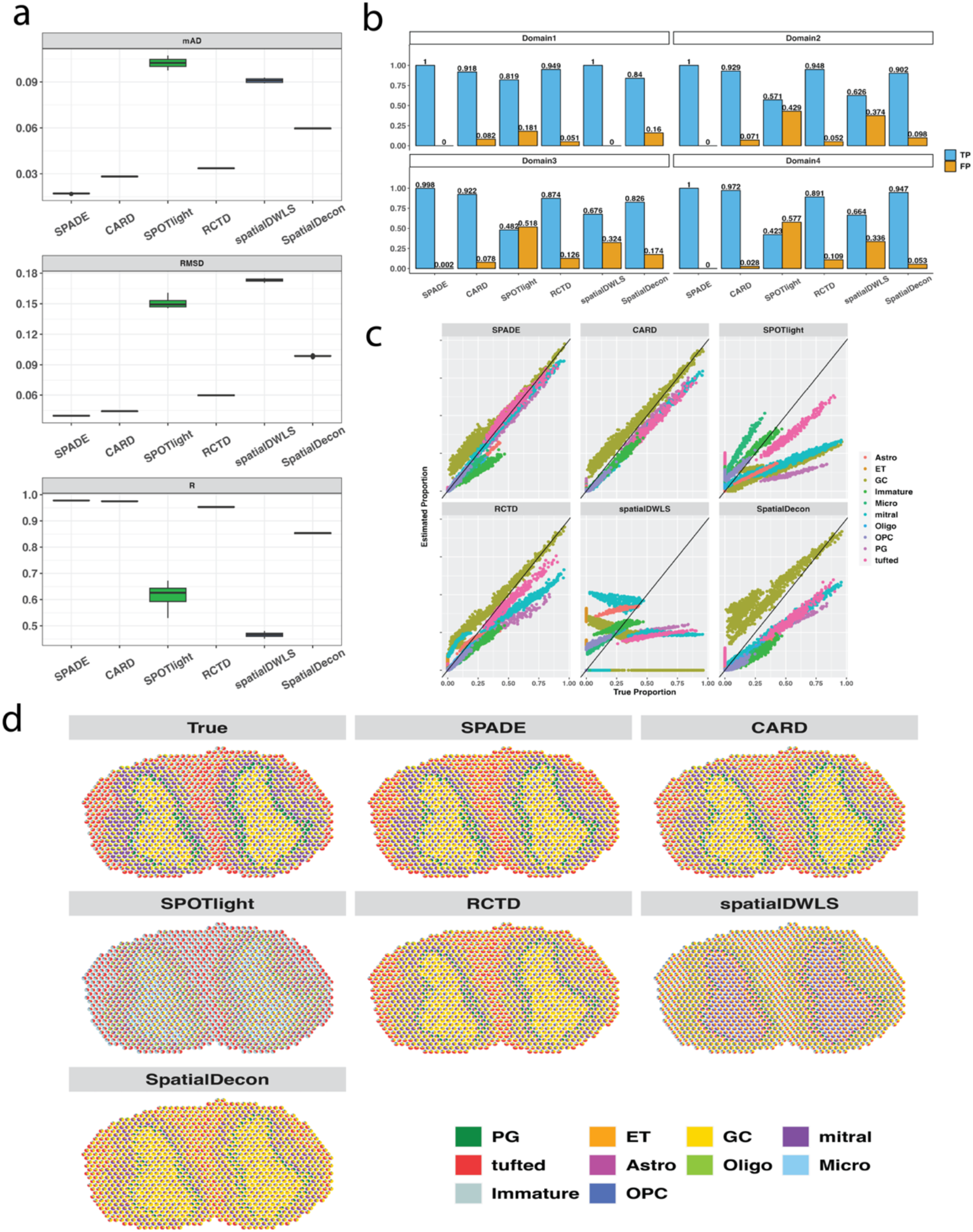
Simulation using mouse olfactory bulb. a. Boxplot for 10 simulations. For mAD and RMSD, the lower value indicates a better performance, while for correlation R, the higher value is preferred. b. True positive and false positive for each domain. c. Dot plot for comparing the estimated proportion with the true proportion in one replicate. The x-axis is the true proportion, y-axis is the estimated proportion. Each dot represents a cell, with a color depicting a cell type. A 45-degree line indicates the same value for true and estimated proportion. d. Spatial scatter pie plot shows estimated cell-type composition on each spatial location from different deconvolution methods, compared to true distribution. Colors represent cell types, and each location is indicated by a pie plot showing the proportions for each cell type.

The dot plot in Figure 2c demonstrated that SPADE achieved the best alignment of cell types to the 45-degree line, indicating that estimated proportions were closer to the true proportion than those by SPOTlight or RCTD. SPADE also has the highest correlation, and the lowest RMSD and mAD across all cell types. When mapping inferred cell type proportions to each spatial location using the spatial scatter plot in Figure 2d, SPADE produced an overall pattern that was highly parallel to the true patterns, outperforming other methods. In conclusion, SPADE is a robust and accurate method for spatial deconvolution, superior to existing methods in both single and multiple simulations.

### Real data application on chicken heart

The heart is the first organ to develop during embryogenesis and interactions among various cell populations play a pivotal role in driving cardiac fate decision. The heterogeneity of cell types in heart development poses a challenge to study by traditional methods. Therefore, it is important to explore new techniques for prediction of cell type heterogeneity during heart development.

Early embryonic development sees the heart start as a simple tube and undergo complex morphological changes to develop into a fully formed four-chambered heart with functional blood vessels. In a previous study, spatially resolved RNA-seq was combined with high-throughput single-cell RNA-seq to investigate the spatiotemporal interactions and regulatory programs driving embryonic chicken heart development [18]. This study utilized chicken embryos to generate over 22,000 single-cell transcriptomes across four key developmental stages, along with spatially resolved RNA-seq on 12 heart tissue sections at the same stages. These stages included day 4, an early stage of chamber formation and the initiation of ventricular septation; day 7, when four-chamber cardiac morphology is initiated; day 10, which is the mid-stage of four-chambered heart development; and day 14, marking the late stage of four-chamber development.

The anatomical development of different regions across four timepoints is demonstrated through H&E stained images (Figure 3a). After applying SPADE, spatial domains were defined for each timepoint, with ventricular separation becoming apparent on Day 4 (Figure 3b). From Day 7 onwards, the clustering of different chambers could be clearly observed. The single-cell RNA-Seq data underwent filtering and preprocessing to select the top variable genes between cell types, which were then aggregated to generate cell type counts for deconvolution. The scatter pie plot (Figure 3c) reveals the predominance of immature myocardial and fibroblast cells on day 4 of heart development, with a decrease trend in the proportion as the hearts mature. This phenomenon is attributed to an initial tube-like structure of the chicken heart during early developmental stages [19], which necessitates the presence and active participation of fibroblast cells in the formation of connective tissue. As heart development progresses, fibroblast cells undergo proliferation and differentiation into various types of connective tissue cells. During later stages of development, the number of fibroblast cells in the heart begins to decline as the heart matures and becomes specialized. However, fibroblast cells continue to play an important role in maintaining the structure and function of the heart throughout the chicken*’*s life [20]–[24]. On the contrary, the number of cardiomyocyte cells significantly increases during the development of the chicken heart, with the highest rate of proliferation occurring from day 4 to day 7, and slowing down from day 10 to day 14.

**Figure 3:**
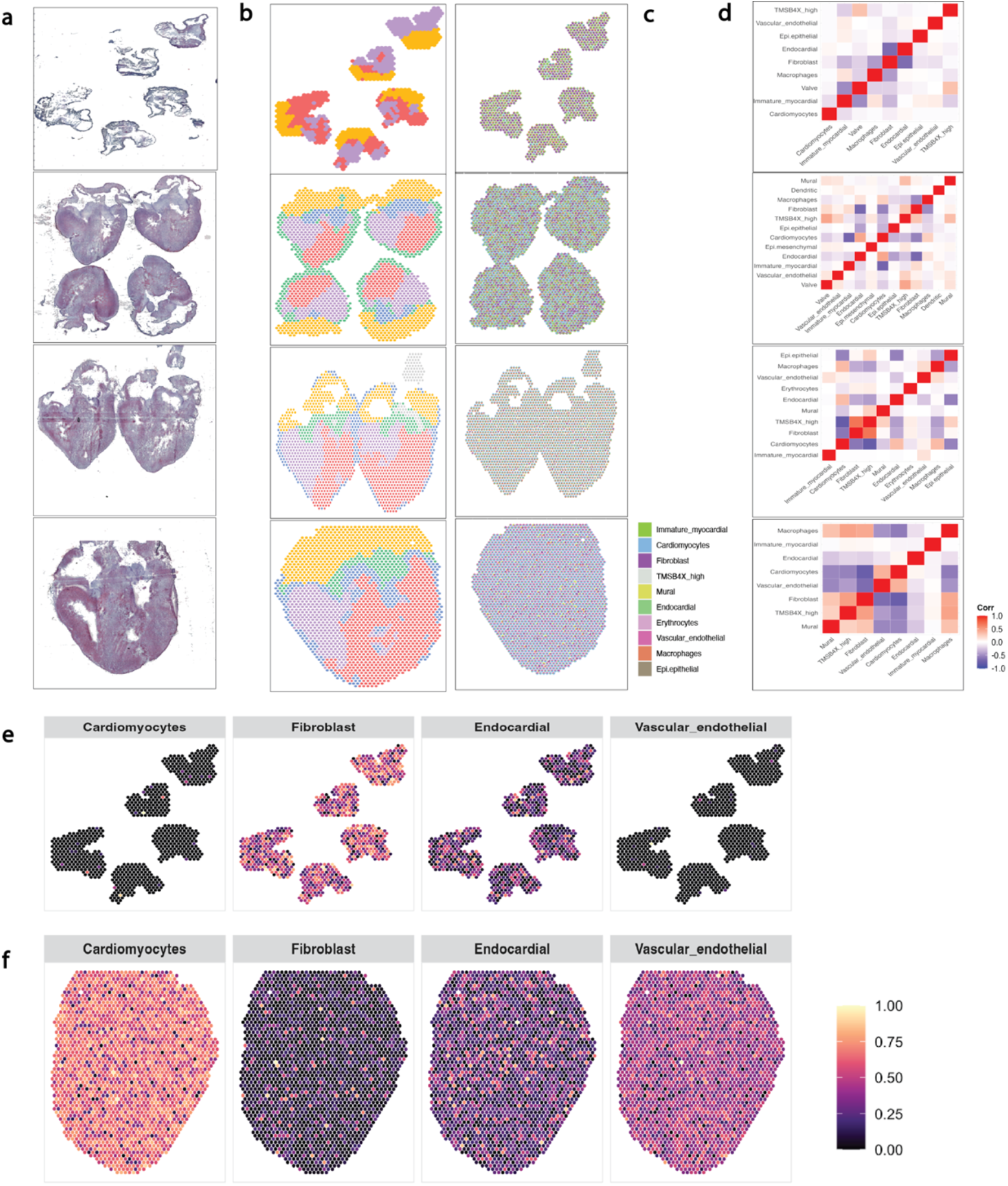
Cell type estimation during chicken heart development. a. H&E stained images for hearts at different days are downloaded from the GitHub [35]. b. Detected spatial domain. Color represents different domain. c. Scatterpie plot for cell type proportion for each day. d. Correlation between estimated cell type proportions across locations. e,f. Cell type proportion in each location at day 4 (e) and day 14 (f).

Cardiomyocytes increase dramatically during embryogenesis partly due to cell proliferation. It has been found that the rate of cardiomyocyte proliferation is the highest during the early stages of heart development and decreases as the heart matures [22], [25]–[28]. This is consistent with our results generated from SPADE. The immature myocardial cells consist a subset of cardiomyocytes that only appear on days 4 to 7 of embryonic development. As the heart continues to develop, these immature cells undergo differentiation to become mature cardiomyocytes. This process of differentiation is critical for the contractile function of the heart [18].

The heart contains multiple cell types, including cardiomyocytes, fibroblasts, endothelial cells, and smooth muscle cells. During heart development, these different cell types interact and form complex networks that are essential for the proper functioning of the heart [29]–[32]. We conducted a cellular colocalization analysis by calculating the correlation between cell types in order to evaluate the spatial colocalization patterns between each cell-type pair. This analysis allows us to assess the extent of cell type colocalization, which refers to the physical proximity and interaction between cell types [30, 31]. We observed that the correlation between cell types increased over the stages of the development, suggesting that the spatial coherence of organization increases during the development, as depicted in Figure 3d. To further illustrate this, we visualized the distribution of various cell types, such as cardiomyocytes, fibroblasts, endocardial cells, and vascular endothelial cells, in each location (Figure 3e). Additionally, we examined the expression patterns of selected marker genes across time (Figure 3f) to highlight the regions where they are active. Our findings suggest that the spatial organization of cell types varies across developmental stages.

## Discussion

Spatial transcriptomics data is a crucial source of information regarding the spatial distribution of gene expression patterns in cells. This technique allows identification of regional differences of gene expression within a tissue, an essential element for understanding the biological significance of the tissue or organ. However, without the knowledge of specific cell types present in each region, current version of spatial genomics can be difficult for data interpretation and for revealing the biological significance.

Cell type deconvolution is a computational methodology that can be used to identify specific cell types present in a tissue sample based on gene expression data. By applying cell type deconvolution to spatial transcriptomics data, we can identify the cell types that are present in each region of a tissue. This can help to contextualize the gene expression data and provide insights into the biological processes that are occurring at the cellular level within the tissue. While several cell type deconvolution methods have been developed for spatial transcriptomics data, most of these methods do not consider the spatial domain structure when predicting the number of cell types. SPADE overcomes such limit and integrates spatial transcriptomic data with single-cell RNA-Seq data to infer the fraction of cell types at each location.

After conducting tests on a synthetic dataset generated from a study on the mouse olfactory bulb, we found that SPADE exhibited exceptional performance. The simulation produced results with remarkable accuracy in predicting both the cell types and the number of cell types present in various locations. Through the application of the SPADE algorithm on chicken heart development data, we acquired insights into the development of specific cell types throughout four stages of ventricular development. These findings provided valuable knowledge basis that can be extrapolated to inform future clinical studies. For example, investigation of human breast cancer enables the discernment of cell type heterogeneity within and between distinct subtypes of breast cancer. Such insight has the potential to guide comparative analyses and enhance our understanding therefore treatment of this complex disease.

The performance of SPADE can be enhanced through incorporation of a better designed reference dataset. In this study, we utilized only one single cell RNA-Seq dataset as reference, potentially limiting the algorithm*’*s overall efficacy. Recent advancements in scRNA-Seq technologies have facilitated the generation of multiple reference datasets from different platforms or samples obtained from the same tissues. The integration of these diverse scRNA-Seq datasets holds the potential to provide a comprehensive and accurate reference set, thereby improving the performance of SPADE.

## Methods

### Spatial domain detection

Spatial domain detection [15], [36], [37] is an essential step for spatial transcriptomics. A spatial domain refers to regions that exhibit spatial coherence in both gene expression and histology. Conventional methods for identifying spatial domains rely on clustering algorithms that take gene expression as the only input and ignore spatial information and histology [15]. To account for spatial correlations, spaGCN integrates gene expression, spatial location, and histology to construct a graph convolutional network for identifying both spatial domains and domain-specific genes. The spaGCN algorithm consists of three steps. First, information from physical location and histology is utilized to construct an undirected graph that reflects the relationships between all spots. Second, a graph convolutional layer is applied to integrate gene expression data from adjacent locations. Lastly, accumulated expression data is grouped by an unsupervised iterative clustering algorithm, with each cluster treated as a spatial domain. Subsequently, domain-specific genes can be identified through differential gene analysis. Further details can be found in the original publication [15].

### Determine the number of cell types for each domain

A crucial disparity between bulk deconvolution and spatial deconvolution is that not all cell types are uniformly distributed across all regions. Thus, accurately determining the presence of specific cell types in each location is imperative for effective cell type deconvolution. Based on the premise that locations within a given domain are closely correlated, it is hypothesized that each domain comprises of same cell types, differing only in their relative proportions. To address this, a lasso-regularized generalized linear model [38] is employed to select cell types for each domain through the following methodology:

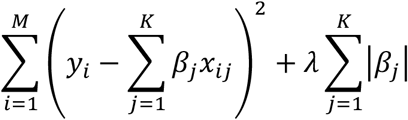

where *y*_*i*_ is the gene expression for gene *i, x*_*ij*_is the gene *i* expression for cell type *j*, β_*j*_is the coefficient for cell type *j*. By estimating the cell type coefficient, we are performing a cell type selection step, if a coefficient shrinks to 0, the corresponding cell type is not present for that location. λ is the tunning parameter and is chosen by 10-fold cross-validation.

Once obtained the cell type related coefficient matrix for each location within each spatial domain, we convert it to a binary matrix, with each entry has a value 1 or 0. Specifically, we implemented an adaptive thresholding that employed a 2D convolution using the Fast Fourier Transform (FTT) to filter the coefficient matrix. This approach enabled us to efficiently identify entries that surpassed a certain threshold. Specifically, if a coefficient exceeded the filtered value, the corresponding entry was set to 1, while entries that fell below the threshold were set to 0.

### Cell type proportion estimation for each location and each domain

The deconvolution problem can be solved to find the optimal estimation for cell type proprotion that minimize the difference between estimated spatial gene expression and observed spatial gene expression for each location as below:

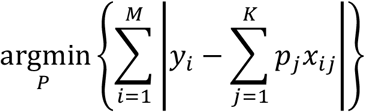

where *y*_*i*_ is the expression for gene *i*(= 1 … *M*). *x*_*ij*_is the expression for gene *i* for cell type *j* (= 1 … *K*) that is extracted from single-cell reference. *p*_*j*_ is the proportion for cell type *j*.

Here we select the absolute deviation loss as the optimal choice due to its less sensitive to the extreme values than the commonly used quadratic loss function. The optimization problem is solved using the Augmented Lagrange Minimization algorithm that is implemented by auglag function in R package alabama. Due to the unique feature of proportion, we not only minimize the nonlinear objective function, but also supply two constrains. Each element of the P matrix has to be nonnegative, and the sum of all cell type within each sample needs to be 1.

### Construct reference

The first step is to generate reference data that is used for inferring gene distribution across cell types. We select single-cell RNA-Seq data that contains tissue or samples that have similar phenotype to the bulk data. Single-cell data have been checked for quality based on the standard pre-processing workflow, including the unique feature count for each cell and the percentage of reads mapped to the mitochondrial genes [42]. The reference basis is built following the main idea from MuSiC [8] which contains the following steps. First, we calculate the cross-cell variation for each gene of each cell type within an individual sample to account for cell type and sample specific library size. In particular, we subset the expression data by removing redundant cell type annotations given by original single-cell study and by removing genes with 0 counts. For each sample within each cell type, we scale the gene expression by their library size, which is calculated by summing all gene counts for each cell. Next, we filter genes by three criteria to keep the genes that satisfy these criteria: 1) the genes shared between single-cell data and bulk data, 2) commonly used cell type biomarkers or highly cited markers, and 3) differentially expressed genes (DEGs) by comparing each pair of cell types. We used a well-known tool Seurat for DEGs detection [43]. The resulting table is a gene by cell type expression matrix that can be implemented in the cell type deconvolution model.

## ACKNOWLEDGEMENTS

This research was partially supported by NIH R01 GM125212, R01 GM126165, and Holsclaw endowment to Q.M.C; NIH 1R01GM139829, 1P01AI148104-01A1, and USDA ARZT-1361620-H22-149 to L.A.

